# Effects of Antipsychotic Drugs and Potassium Channel Modulators on Cognition-related Local Field Potential Spectral Properties in Mouse Hippocampus and Frontal Cortex

**DOI:** 10.1101/2020.05.26.116343

**Authors:** Dechuan Sun, Mojtaba Kermani, Matt Hudson, Xin He, Ranjith Rajasekharan Unnithan, Chris French

**Affiliations:** Department of Medicine, The University of Melbourne; Department of Electrical and Electronic Engineering, The University of Melbourne; School of Biomedical Sciences, Monash University; Department of Neuroscience, Monash University

## Abstract

Local field potentials (LFPs) recorded intracranially display a range of location specific oscillatory spectra which have been related to cognitive processes. Although the exact mechanisms producing LFPs are not completely understood, it is likely that voltage-gated ion channels which produce action potentials and patterned discharges play a significant role. It is also known that antipsychotic drugs (APDs) affect LFPs spectra and a direct inhibitory effect on voltage-gated potassium (*K*_*v*_) channels has been reported. Additionally, *K*_*v*_ channels have been implicated in the pathophysiology of schizophrenia, a disorder for which APDs are primary therapies. In this study we sought to: i) better characterise the effects of two APDs on LFPs and connectivity measures and ii) examine the effects of potassium channel modulators on LFPs and potential overlap of effects with APDs. Intracranial electrodes were implanted in the hippocampus (HIP) and pre-frontal cortex (PFC) of C57BL/6 mice; power spectra, coherence and phase-amplitude cross frequency coupling were measured. Drugs tested were the APDs haloperidol and clozapine as well as voltage-gated potassium channel modulators (KVMs) 4-aminopyridine(4AP), tetraethylammonium (TEA), E-4031 and retigabine. All drugs and vehicle controls were administered intraperitoneally. Both APDs and KVMs significantly reduced gamma power with the exception of 4AP, which conversely increased slow-gamma power. Clozapine and retigabine additionally reduced coherence between HIP and PFC. Phase-amplitude coupling between theta and gamma oscillations in HIP was significantly reduced by the administration of haloperidol and retigabine. These results provide previously undescribed effects of APDs on LFP properties and demonstrate novel modulation of LFP characteristics by KVMs that intriguingly overlaps with the effects of APDs. The possibility of a common mechanism of action deserves further study.

## Introduction

APDs are commonly used for the treatment of psychotic illness. Considerable interest has developed around the hypothesis that some aspects of psychosis may be related to impairment of intrinsic cerebral rhythmicity detectable in the EEG as well as LFPs in subregions of the brain (Lee et al., 2003). The observation that APDs can affect EEG and LFPs properties raises the possibility of a causal therapeutic connection and also illustrates the need for better understanding of the genesis of these oscillations (Moran and Hong, 2011; Jackson and Severiratnel, 2019). While synaptic currents are undoubtedly significant in EEG and LFPs generation, it has also been recognised that voltage-gated ion channels underlie neuronal rhythmicity (Wang, 2010), but the effects of voltage-gated ion channel modulation of *in vivo* oscillations have not been previously explored. We therefore firstly sought to extend observations of the effects of APDs clozapine and haloperidol on LFPs spectra in brain regions involved in psychotic illness (Muller, 1985; Ghoshal and Conn, 2015; Lieberman et al., 2018) by measuring coherence as well as cross frequency coupling. These phenomena have been proposed to be measures of functional connectivity between and within brain regions (Fries, 2015; Salimpour and Anderson, 2019), and impaired connectivity has been reported in psychotic illness studies.

APDs have been shown to directly inhibit neuronal potassium channels at therapeutically relevant concentrations (Ogata et al., 1985; Nakazawa et al., 1995). In view of this, we hypothesised there might be an overlap of effects with APDs, so the same observations were made with a range of KVMs for different *K*_*v*_ channel subtypes: – tetraethlammonium (TEA) known to affect non-inactivating “delayed rectifier” currents, 4-aminopyridine (4AP) blocks inactivating “A type” potassium currents, the ERG channel blocker E4031 and a *K*_*v*_7 channel activator retigabine.

## Methods

### Animals

Fifty-six adult male C57BL/6 mice (aged 6-10 weeks, 22-26g) were group-housed (4 mice/cage) in the Biological Research Facility of the Department of Medicine, Royal Melbourne Hospital, University of Melbourne. The mouse housing room was maintained in a 12-h light/dark cycle (lights on from 7:00 am to 7:00 pm) with autoclaved water and rodent laboratory chow ad *libitum*. The mice were handled for 10 minutes each day during the daytime for seven days before the commencement of experiments. All surgical and experimental procedures were approved by the University of Melbourne/Florey Animal Ethics Committee (AEC No. 16-019UM).

### Drugs

E-4031 (Alomone-Labs, IL) and TEA (Sigma-Aldrich, USA) were dissolved in 0.9% saline. 4-AP (Sigma-Aldrich, USA) and haloperidol (Sigma-Aldrich, USA) were dissolved in a vehicle solution of 8% dimethyl sulfoxide (DMSO) and 92% saline (0.9%). Clozapine (Sigma-Aldrich, USA) was dissolved in pure acetic acid, then diluted with saline (0.9%) to make a 0.1% acetic acid solution. Retigabine (Leader-Biochemical Group, CHN) was dissolved at 37°C in a vehicle solution of 6% Tween 80 and 94% saline (0.9%).

### Electrode implantation surgery

Mice were anesthetized with isoflurane and oxygen (induction: 3%, maintenance: 1%-2.5%), placed on a heat pad (37°C), fixed in a mouse stereotaxic frame, and subcutaneously injected with the analgesic carprofen (Rimadyl^®^,Pfizer Animal Health) at 1:100 dilution (0.01 ml/g). The head was shaved and sterilized with betadine and ethanol. A single incision along the scalp midline was made on the scalp, connective tissue was removed and hydrogen peroxide (Sigma-Aldrich, USA) was applied to clean the skull surface. After aligning the skull, two custom length (10 mm, 13.5 mm) tungsten electrodes (diameter:270um; Ginder Scientific, CAN) were implanted into the CA1 region of the hippocampus (AP −1.8 mm, ML 1.3 mm, DV −1.4 mm) and PFC (AP 2.0 mm, ML 0.3 mm, DV −1.7mm), and a reference/ground electrode was placed 2 mm posterior and 2 mm unilateral to lambda, overlying the cerebellum. The electrodes were connected to a multi-channel socket connector (Harwin, AU) with 1mm heat shrink on the outside of the middle pin to adequately isolate the electrodes. Two additional 1mm screws were implanted (±AP 1.8 mm, ML −1.6 mm) for mechanical stabilisation. Cyanoacrylate and dental cement were then carefully applied to secure the electrodes, anchors and connector together. The mice received an additional intra-peritoneal (i.p.) injection of carprofen 24 hours following the surgery and allowed 8 days recovery before commencement of further procedures.

### Experimental protocol

All experiments were performed at 8:00 pm (AECST). On the day of experiment, the mice were brought into a silent recording room 30 min before the start of recording to acclimatise. At the start of each session, the mouse was placed into a Lucite restrainer (80 mm wide, 200 mm long) within a Faraday cage and the electrode connector attached to a custom-made, light-weight, three-channel shielded cable. Each recording session began with a 10 min habituation period, followed by a 15 min baseline EEG recording. The mouse was then injected (i.p.) with drug and returned to the restrainer without delay for another 30 min. Seven cohorts (n=7*8) of animals were used. Each animal in the first 6 cohorts received three doses of one drug with 3 days in between sessions. The last cohort of animals received vehicle control solutions. Eight animals were excluded from the analysis due to either poor-quality signal or head-piece dysfunction. Drug doses and animal numbers were as follows:

Doses of drug: haloperidol (0.1 mg/kg, 0.3 mg/kg, 0.9 mg/kg, n=6), 4-AP (0.5 mg/kg, 1 mg/kg, 2 mg/kg, n=7), TEA (0.1 mg/kg, 0.3 mg/kg, 0.5 mg/kg, n=7), retigabine (5 mg/kg, 12.5 mg/kg, 20 mg/kg, n=6), clozapine (0.4 mg/kg, 1.2 mg/kg, 3.6 mg/kg, n=7), E-4031 (10 mg/kg, 20 mg/kg, 30 mg/kg, n=7), vehicle(n=7). The drug doses were derived from previous studies (Aceto et al., 1969; Yamaguchi et al., 1992; Hirano et al., 2007; Kurokawa et al., 2015; Hudson et al., 2016).

### Data acquisition processing

LFP signals were recorded continuously at 20KHz with a 32-channel amplifier (Intan Technologies, RHD2132 amplifier board with RHD2000 USB Interface Board), with the on-board analogue bandpass filter set to be 1-150 Hz. The average power spectra of LFP oscillations in different brain regions (HIP and PFC) and the amplitude coherence between them were calculated using the Chronux package for MATLAB (The MathWorks, USA) (Bokil et al., 2010). The phase-amplitude cross frequency coupling was evaluated with the Fieldtrip toolbox for MATLAB (Oostenveld et al., 2011). For the power spectra and coherence analysis, the signals were first down-sampled to 2000 HZ, segmented into 2s epochs, detrended, and band-pass filtered (2-100 Hz) with a zero-phase filter. The epochs containing movement artefact were deleted from the analysis with a custom-made Matlab script. Continuous multi-taper-based power spectrum and coherence (Mitra and Pesaran, 1999) were measured in each epoch with the time-bandwidth product of 3 and five leading tapers at four frequency intervals: theta (>4-12 Hz), beta (>12-32 Hz), slow-gamma (>32-60 Hz), fast-gamma (>60-100 Hz). And then the average derived for both the baseline and post-injection period. For the phase-amplitude coupling analysis, the same pre-processing method was applied except the epoch length was set to be 10s. The modulation index (Caixeta et al., 2013) was measured in each epoch and then averaged in both the baseline and post-injection period to quantify the strength of cross-frequency coupling. The time window used to calculate the complex power spectra was 4 cycles long and, therefore, differed over frequencies, and the number of phase bins was set to be 20. For each animal, the first 15 min baseline (pre-injection) recording served as a self-control and the post-injection response was then expressed as a percentage change with respect to the baseline value.

### Statistical analyses

One-way analysis of variance (ANOVA) was used for all outcomes, and Bonferroni’s post hoc analysis was applied where appropriate. All results are expressed as means ± standard error of mean (S.E.M) or as a percentage variation with respect to the baseline value. Statistical comparisons were performed using SPSS (IBM, Armonk, NY, United States). Statistical significance was set at p<0.05 for all analyses.

## Results

### Drug effects on hippocampal and pre-frontal cortex power spectra

All drugs had significant effects on power spectra of LFPs in the hippocampus in one or more bands. One-way ANOVA displayed significant differences between treatment groups [theta: F_18,126_ =3.465, P<0.001; beta: F_18,126_ = 4.994, P<0.001; slow-gamma: F_18,126_ =46.661, P<0.001; fast-gamma: F_18,126_ =23.720, P<0.001]. The vehicle group remained relatively stable before and after injection. Post hoc comparison identified robust effects of clozapine (3.6 mg/kg, 1.2 mg/kg) and haloperidol (0.9 mg/kg, 0.3 mg/kg) on reducing slow-gamma power with maximal reductions of 31.16% ± 2.84% and 25.33% ± 3.02% compared with baseline (Figure 1A). In the fast-gamma band, all doses of haloperidol significantly decreased power spectra power, with a maximal power reduction of 39.52% ± 4.13% (0.9 mg/kg), while clozapine (3.6 mg/kg, 1.2 mg/kg), produced a reduction of 45.52% ± 4.09% and 44.31% ± 4.20%. Retigabine (20 mg/kg, 12.5 mg/kg) showed the strongest attenuation of both slow-gamma and fast-gamma power, with a maximal reduction of 42.62% ± 3.83% and 63.07% ± 3.32% respectively. TEA (0. 5 mg/kg) was the only drug tested that significantly affected theta band power with a 36.52% ± 7.59% reduction (Figure 1A). It also significantly reduced beta (38.42% ± 4.34%), slow-gamma (23.77% ± 1.90%) and fast-gamma (46.03% ± 4.08%) power. The ERG channel blocker E4031(30 mg/kg) suppressed slow-gamma (23.92% ± 3.64%) and fast-gamma (38.60% ± 4.15%) power (Figure 1A, Figure 2A). 4-AP demonstrated the opposite effect, significantly enhancing slow-gamma power (2 mg/kg,56.67% ± 6.65%; 1 mg/kg, 15.52% ± 2.93%), but had no clear effects on fast-gamma and theta power.

**Figure 1.**
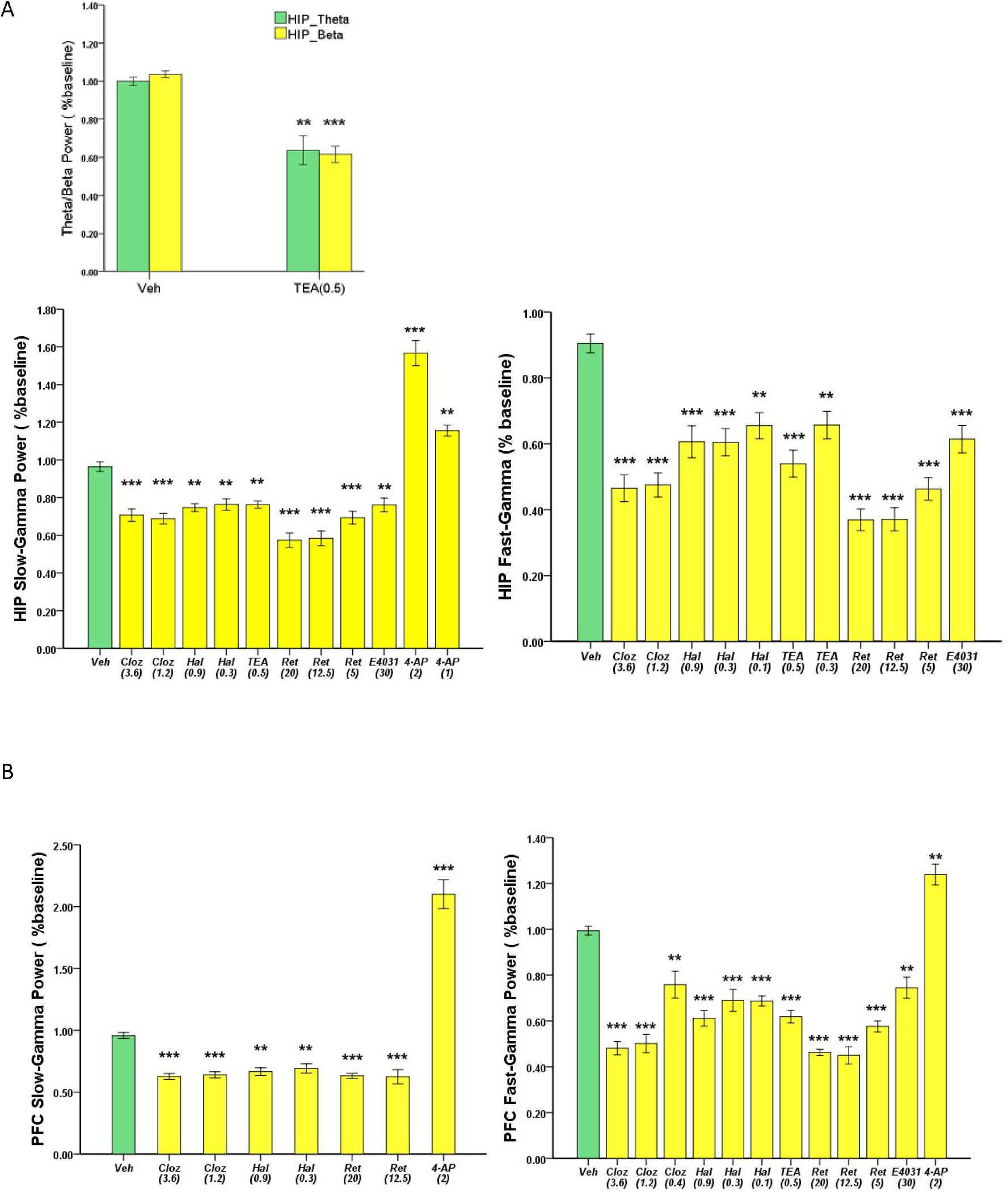
Effects of drugs on the power of different frequency bands in (A) HIP (B) and PFC (mean ± S.E.M), only significant results are plotted. Clozapine (Cloz), haloperidol (Hal), TEA, retigabine (Ret), E-4031 showed suppressed power, mostly in the gamma band, whereas 4-AP produced enhancement. The numbers in the parentheses under drug names indicate dose, with unit mg/kg. ***p<0.001, **p<0.01, *p<0.05 displayed the significant difference between vehicle and drugs.

**Figure 2.**
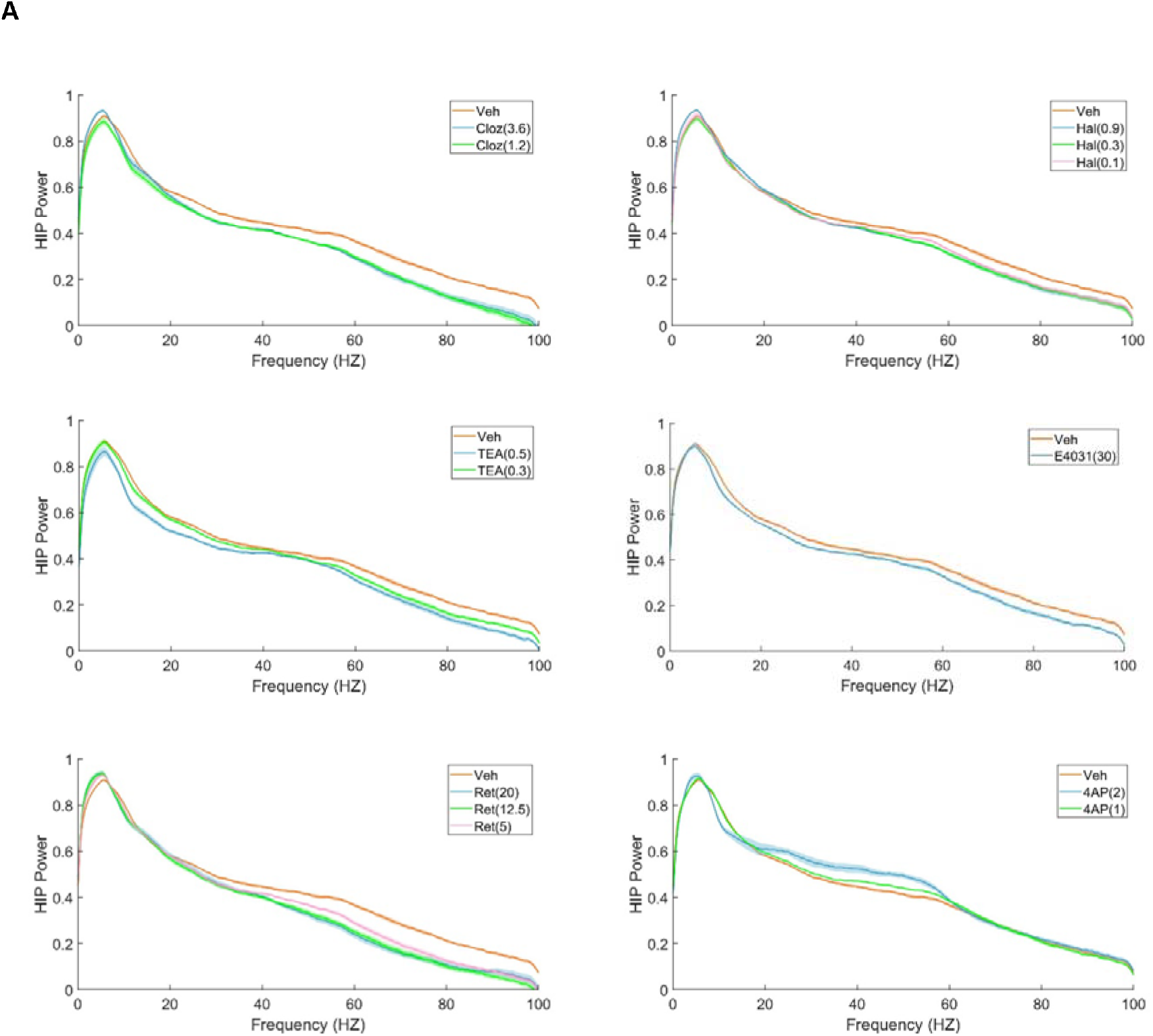
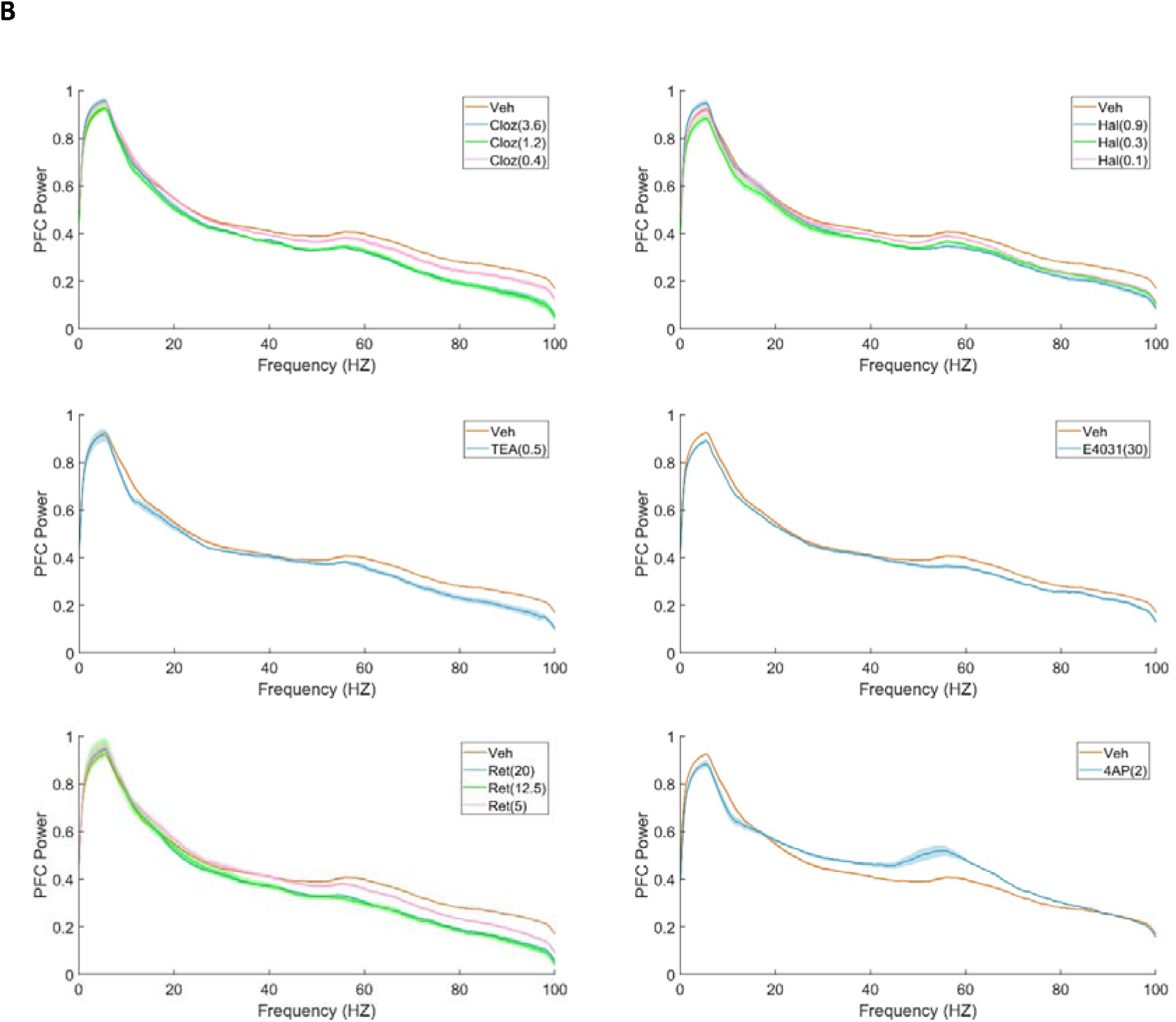
Averaged power spectrum after administration of vehicle and different drugs in (A) HIP and (B) PFC. The data are log-transformed to better show the difference. Only significant results are displayed.

A similar pattern was seen in the prefrontal cortex recordings with significant differences observed between treatment groups [slow-gamma: F_18,126_ = 56.022, P<0.001; fast-gamma: F_18,126_ = 39.552, P<0.001]. Post hoc analysis showed that both clozapine (3.6 mg/kg, 1.2 mg/kg) and haloperidol (0.9 mg/kg, 0.3 mg/kg) significantly decreased slow-gamma power, with a maximal reduction of 37.26% ± 2.44% and 33.43% ± 3.09% respectively. All three doses of both APDs showed significant suppression of fast-gamma power [clozapine: 51.91% ± 2.92% (3.6 mg/kg), 49.86% ± 4.01% (1.2 mg/kg), 24.17% ± 5.84% (0.4 mg/kg); haloperidol: 38.86% ± 3.43% (0.9 mg/kg), 31.00% ± 4.81% (0.3 mg/kg), 31.31% ± 2.19% (0.1 mg/kg)]. The *K*_*v*_ activator retigabine (20 mg/kg, 12.5 mg/kg) was again the most potent drug to attenuate slow-gamma (37.46% ± 2.18%) and fast-gamma (54.99% ± 3.78%) oscillations to almost the same extent individually. Interestingly, TEA did not impact slow-gamma power significantly, but did reduce fast gamma power by 27.46% ± 1.91% at the doses of 0.5mg/kg. The ERG channel blocker E-4031 produced a 25.53% ± 4.65% reduction in fast gamma power, but not in the slow-gamma band.

Contrary to the attenuation effects of the other KVMs, 4-AP at doses of 2mg/kg very strongly increased slow-gamma power (110% ± 11.64%), and also increased fast-gamma power (23.90% ± 4.51%), different from the response seen in hippocampus (Figure 1B, Figure 2B). All the corresponding p values were summarized and described in Figure 1.

### Effects of Antipsychotic Drugs and Potassium Channel Modulators on Hippocampal and Prefrontal Cortex Coherence

Correlation in the local field potential recorded from two brain areas is often characterized by computing the coherence, which reflects the degree of phase consistency across trials between two sites (Srinath et al., 2014). It is considered likely to be a measure of information flow and storage (Siapas et al., 2005), and takes values from 0 to 1. Significant group treatment differences were observed after applying one-way ANOVA [theta: F_18,126_ =6.700, P<0.001; fast-gamma: F_18,126_ =6.329, P<0.001]. Post hoc analysis revealed that clozapine (3.6 mg/kg, 1.2mg/kg) and retigabine (20 mg/kg, 12.5mg/kg) were the only two compounds that affected coherence between HIP and PFC, with maximal reductions of 28.20% ± 3.03% and 23.49% ± 2.73% respectively in the theta band, and 23.42% ± 6.39% and 25.93% ± 3.58% in fast-gamma band with respect to the baseline (Figure 3). All the corresponding p values were summarized and described in Figure 3.

**Figure 3.**
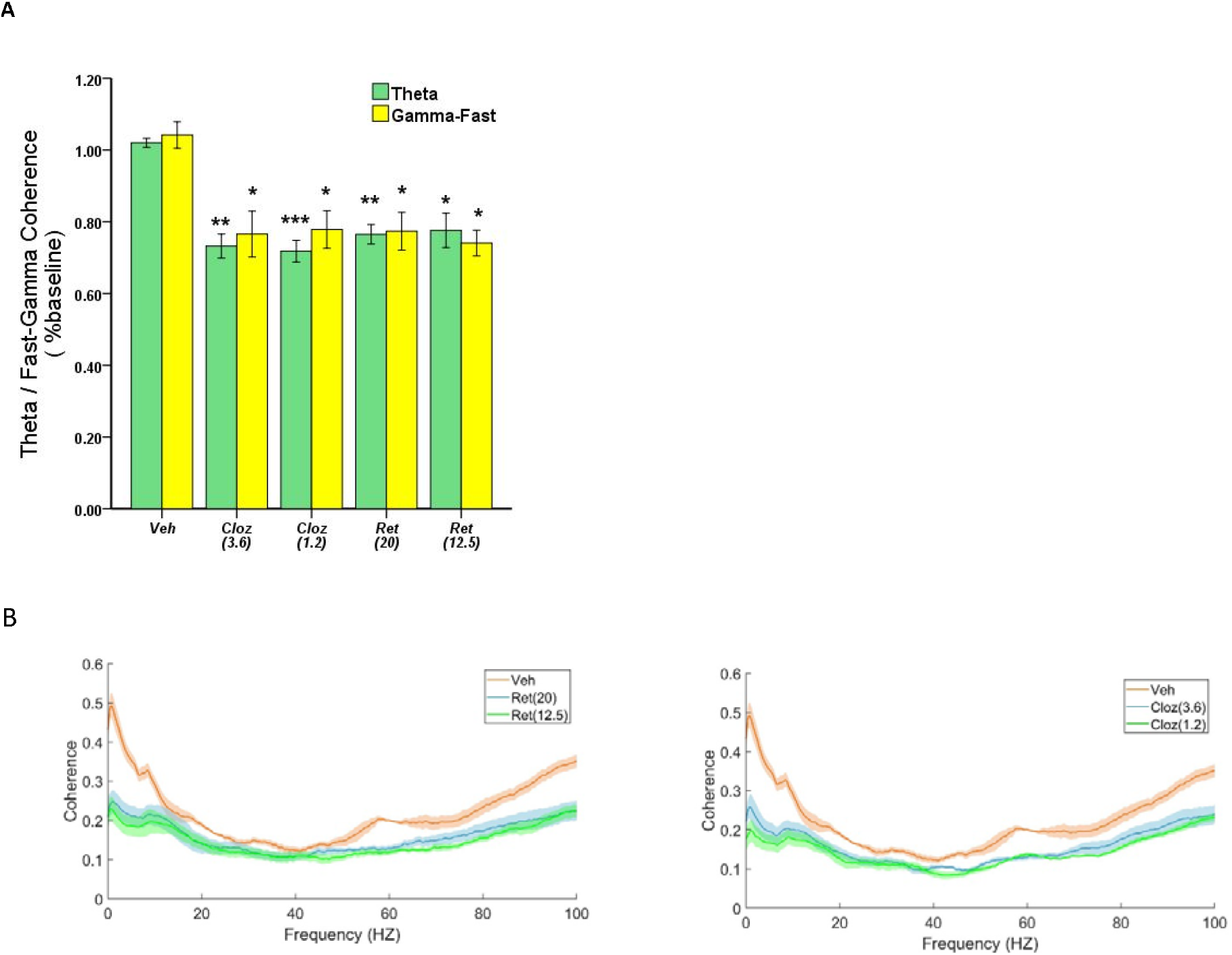
(A) Effects of vehicle, clozapine and retigabine on HIP-PFC coherence in the theta and fast-gamma frequency bands. (B) Average results after administration of vehicle and drugs. ***p<0.001, **p<0.01, *p<0.05

### Effects of Drugs on Cross Frequency Coupling in the Hippocampus and Prefrontal Cortex

Cross-frequency coupling refers to the phase and spectral components recorded from a single site and their relative relationship. Brain rhythms at different frequencies appear to interact, and this can be quantified as a “modulation index”. It has been hypothesised that the phase of low frequency signals (such as theta band) can modulate the amplitude of high frequency components including the gamma band. It has been suggested that cross-frequency coupling (CFC) might play a functional role in neuronal computation, communication and learning (Canolty and Knight, 2010). We analysed the phase-amplitude coupling in both HIP and PFC between theta and gamma bands. For this purpose, the slow and fast gamma range were combined. Significant CFC was not observed within PFC, but one-way ANOVA revealed significant treatment groups differences [F_18,126_ =5.990, P<0.001] within HIP. Significantly decreased modulation indexes between theta band and gamma band were observed in both haloperidol (0.9 mg/kg, 8.52% ± 2.11%, p<0.01) and retigabine (20 mg/kg, 8.50% ± 1.65%, p<0.01; 12.5mg/kg, 8.75% ± 2.43%, p<0.01) groups compared with vehicle (0.04% ± 0.66%). No consistent effects were seen with other drugs (Figure 4A, Figure 4B).

**Figure 4.**
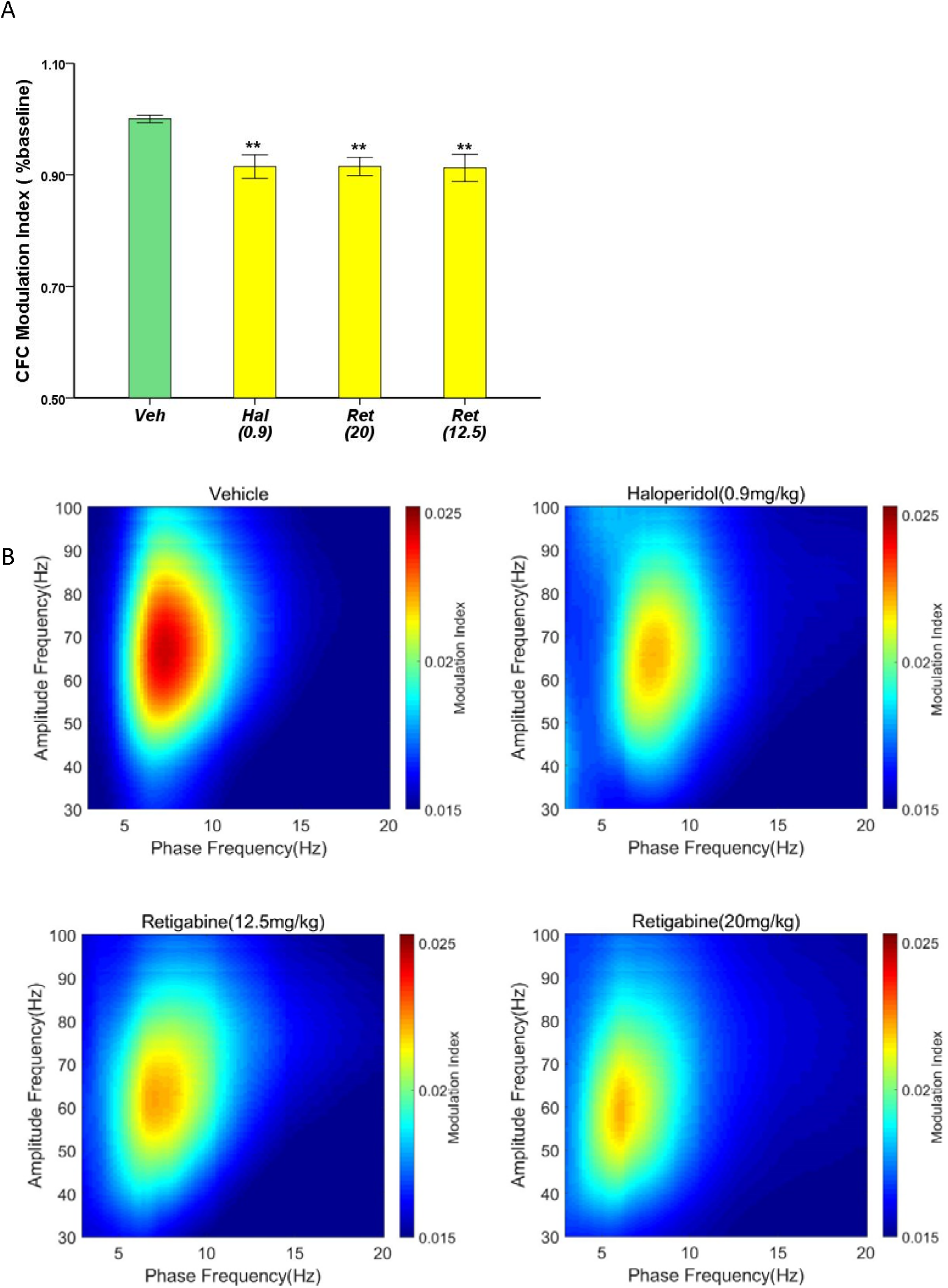
(A) The effects of vehicle, haloperidol and retigabine on the phase-amplitude coupling between theta and gamma bands. (B) The average modulation index after the administration of vehicle and drugs [haloperidol (0.9 mg/kg), retigabine (20 mg/kg), retigabine (12.5 mg/kg) respectively]. Compared with the vehicle group, significant reductions of modulation index were not observed in the other drug groups. **p<0.01 displayed the significant difference between vehicle and drugs.

## Discussion

Low amplitude voltage oscillations were first observed with scalp recorded human brain activity by Berger (1929). Abnormalities of these signals were soon associated with epilepsy and more recently disorders associated with cognitive dysfunction such as schizophrenia and Alzheimer’s disease manifest EEG changes (Uhlhaas and Singer 2015; Cassani et al., 2018). It remains uncertain to what extent these rhythms are simply the result of ensemble synaptic and neural spiking activity, or alternatively, are mechanistically significant processes. Nevertheless, evidence is accumulating that these rhythms have intrinsic functional roles, such as ensemble synchronisation for memory encoding and communication across brain regions (Colgin, 2013).

Individuals with psychotic disorders, even naive patients, who display abnormalities in EEG high frequency spectra could get certain improvement with the treatment of APDs (Won et al., 2018), and APDs also affect *in vitro* (Gao 2007; Jones et al., 2012). We therefore sought firstly to confirm previous observations of EEG changes in response to APDs, in particular haloperidol (termed as “typical” antipsychotic) and clozapine (classified as “atypical”). This nomenclature relates to the earlier discovered “typical” APDs with strong dopamine 2 receptor antagonism and a tendency to cause extrapyramidal side effects such as tardive dyskinesias. Later developed “atypical” drugs tend to have significant 5HT receptor interactions, less extrapyramidal side effects and clinically appear to have some ameliorative effect on the so-called negative symptoms including cognitive impairment (Leucht et al., 2009; Fusar-Poli et al., 2014). We also sought to extend these observations by measuring coherence which may be a marker of inter-regional information integration, as well as cross frequency coupling, which has been suggested to be a mechanism for effective local information processing (Jensen and Colgin 2007; Canolty and Knight, 2010). We additionally measured the effects of potassium channel modulators on these EEG properties for several reasons. Firstly, in addition to synaptic conductance, voltage-gated ion channels, including potassium channels, are predicted to have a significant role in the intrinsic rhythmicity of neural networks, especially theta and gamma ranges (Wang, 2010). APDs have also been reported to directly inhibit various types of potassium channels at therapeutically relevant concentrations in neuroblastoma and pheochromocytoma cells (Ogata et al., 1985; Nakazawa et al., 1995), observations we have replicated in rat hippocampal neurons (Chan and French, 2011). Antipsychotics are also known to inhibit *K*_*v*_ 11 subtypes (HERG and ERG) channels which have been implicated in schizophrenia (Shepard et al., 2007; Huffaker et al., 2009) and are expressed widely throughout the rodent and human brain (Wymore et al., 1997). We therefore sought to test the hypothesis that there could be overlap of effects on LFP properties between APDs and KVMs, and this was indeed confirmed.

Both haloperidol and clozapine reduced slow and fast gamma power in the hippocampus and PFC, with no effect on peak frequency. This has been observed in several previous studies (Jones et al., 2012; Hiyoshi et al., 2014; Hudson et al., 2016). We additionally tested for changes in coherence and CFC and found that clozapine, but not haloperidol, reduced coherence between the hippocampus and PFC, perhaps suggesting a functional difference of effect and potential biomarker. It would therefore be of interest to repeat these experiments with other atypical APDs to see if there are similar effects on CFC. Interestingly, there is evidence that clozapine has overall higher efficacy than first generation antipsychotics on negative symptoms compared with haloperidol, although the exact magnitude is still being clarified. Haloperidol was additionally found to decrease the modulation index between theta and gamma bands in the hippocampus.

Several forms of potassium channel modulators were tested, all of which produced effects on LFP spectra. We are unaware of previous studies *in vivo* of LFP or EEG properties of these drugs, although *in vitro* changes in gamma power with Kir3 potassium channel activation have been demonstrated (Johnston et al., 2014) and Leao et al. (2009) have shown effects on spike timing phase relationships with gamma spectra with Kv7 modulation.

TEA is a classical inhibitor of “delayed rectifier” class potassium currents, that is, relatively slowly activating, non-inactivating conductance first identified by Hodgkin and Huxley (1952) in the squid axon. TEA (0. 5 mg/kg) significantly decreased theta power by 36.52%, beta by 38.42%, slow-gamma (23.77%) and fast-gamma power (46.03%) in the hippocampus, but only fast gamma in the PFC. Intriguingly, 4AP had quite different effects compared to TEA, greatly enhancing slow-gamma power (2 mg/kg, 56.67%; 1 mg/kg, 15.52%) in the hippocampus, and showing very large enhancement of the slow-gamma band power in PFC. No behavioural or electrophysiological evidence of epileptic activity was observed with these doses. Given the somewhat different kinetic properties of type A-current (I_A_) and delayed rectifier K-current (I_KDR_), it is not surprising that different effects on LFPs were seen. Differential effects of TEA and 4AP on action potential firing in neural tissue have been demonstrated. In particular, 4AP is important for producing repetitive firing (Connor and Stevens, 1971) with blockade of I_A_ resulting in hyperexcitability of neurons (Segal et al., 1984). It is also ossible that TEA may cause cell depolarisation as has been seen in the squid axon (Wong and Binstock, 1980) which could either increase excitability by lowering the spike threshold, or even reduce excitability by inactivation of sodium currents. Of considerable interest is the recent finding that modified forms of 4-AP enhance cognitive performance in multiple sclerosis patients (De Bakirtzis et al., 2018; Giglio et al., 2019). It would be of interest to determine whether this drug may have cognition enhancing (nootropic) effects in the behaviour of normal animals such as has been demonstrated for caffeine and nicotine (Angelucci et al., 1999; Gould et al., 1999), and conversely whether these and other nootropics might have similar effects to 4AP on LFP properties, potentially providing a useful biomarker.

The antiepileptic drug retigabine was tested as its most well-described effect is to activate KCNQ (M-channel) currents. It enhances the open probability of the channels and increases the slowly activating potassium current I_M_ by hyperpolarising the activation curve and reducing its deactivation, resulting in neuronal hyperpolarization and decreased firing frequency (Tatulian et al., 2001). It might have been expected to have opposite outcomes compared to TEA, but rather had complex effects, showing on the one hand strong attenuation of both slow and fast gamma power in the hippocampus and PFC (similar to TEA) but on the other hand did not reduce beta and theta power in the hippocampus as seen with TEA. Retigabine is also known to inhibit other Kv channel types, which might be related to these effects (Stas et al., 2016). Additionally, Johnston et al. (2014) have demonstrated a reduction in gamma band power *in vitro* using a G-protein coupled potassium channel, Kir3 activator.

The ERG channel antagonist E4031 was tested as these channels have been detected in the rat hippocampus and cortex (Papa et al., 2003). Cardiac tissue is known to be important in heart rhythm generation (Vandenberg et al., 2012). While there are few electrophysiological studies in the central nervous system, Cui et al. (2018) have recently observed effects on neuronal repetitive firing in rat cortical neurons and there has been speculation that ERG family channels may be clinically significant targets of APDs (Kongsamut et al., 2002) as well as possible causal involvement in some forms of schizophrenia (Huffaker et al., 2009). Given the known rhythmogenic effects of these channels in cardiac tissue and the results reported by Cui et al. (2018), we hypothesised that ERG blockade would also affect LFP spectral properties, as was demonstrated, with inhibition of slow and fast gamma (23.92% and 38.60% respectively) in the hippocampus and fast gamma in the PFC (25.53%). This result is discordant with Fano et al. (2012) who found no effect of E4031 on acetylcholine induced gamma activity in brain slices, but this may be related to the method of induction of gamma, as well as the inevitable functional disconnection involved with the brain slice technique.

Although all KVMs affected LFP spectra, there may be other explanations for these effects apart from *K*_*v*_ channel modulation. 4AP has been shown to influence synaptic transmission and may target calcium channels (Gu et al., 2004). TEA has also been shown to influence synaptic plasticity (Pelletier and Hablitz, 1996). Because of the common effect of attenuating gamma frequencies of TEA, E4031 and retigabine, we propose a direct effect on *K*_*v*_ channels. Similarly, the enhanced gamma spectra by 4AP is plausibly ascribed to I_A_ inhibition given the strong direct enhancement of excitability observed after I_A_ reduction with 4AP (Segal et al., 1984; Williams and Hablitz, 2015). This could be further investigated with conductance-based computational models of cortical microcircuits where these mechanisms could be tested in silico. A more exact method would be to perform *in vivo* whole-cell patch clamp recordings to directly measure potassium currents during drug infusions.

Some studies propose that intrinsic brain rhythms may facilitate transfer of information between brain regions (Colgin, 2013) and LFP coherence is likely a measure of such connectivity. It has been found that HIP and PFC coherence increased with learning (Benchenane et al.,2010), is impaired in a mouse model of schizophrenia (Sigurdsson et al.,2010), and that attenuated deficits in a mouse model of schizophrenia correlated with improved HIP and PFC coherence (Bygrave et al., 2019). We therefore firstly observed the effect of antipsychotics on HIP/PFC coherence, which according to our knowledge, does not appear to have been examined previously. Clozapine reduced theta and gamma coherence but not haloperidol. Any implication of functional significance of this result would be speculative, but it would be of interest to see if similar findings occur for other typical and atypical antipsychotics. Retigabine had a similar effect but none of the other KVMs. It would also be of interest to see if links between cognitive events and coherence (Jones and Wilson, 2005) were affected by these drugs specifically.

Cross-frequency coupling has been identified as a potential biomarker for several cognitive processes including neuronal computation, communication and learning (Canolty and Knight, 2010). Therefore, it was of interest to measure the effects of APDs and KVMs given the effects of these drugs on intrinsic rhythmicity and coherence as shown above. Investigations of CFC have been performed primarily in memory related studies and increased CFC may be derived from the enhanced synchronization of the neural networks (see Salimpour and Anderson, 2019 for review). Haloperidol and retigabine decreased the hippocampal modulation index between theta and gamma. Dzyubenko et al. (2017) studied the effects of the first-generation and second-generation APDs haloperidol and olanzapine on a model of hippocampal neuronal networks *in vitro*. Increased overall activity and synchronization was observed when treated with olanzapine, whereas haloperidol had the opposite effect. It would be interesting to investigate if drug-specific effects on behavioural correlated CFC modulation (as described by Tort et al. 2009) could be observed.

In summary, the results of the present study extended previous observations of the effects of APDs on the LFP properties and showed that most KVMs produce very similar effects to APDs. Both haloperidol and clozapine suppressed slow-gamma and fast-gamma spectral power, which intriguingly overlap the effects of different KVMs (TEA, E4031, retigabine). Given the overlap of effects, the mechanisms by which APDs affect LFP spectra may be at least partly due to potassium channel modulation. One exception is 4AP, which displayed the opposite effect on gamma power suppression, and this may be due to its effects on I_A_ inhibition and the different dynamical effects on neuronal firing demonstrated *in vitro*. The reduced PFC/HIP coherence caused by clozapine and retigabine, as well as the deficit observed in hippocampal modulation index caused by haloperidol and retigabine revealed potentially differential modulatory mechanisms of typical and atypical APDs, and this also verified the overlap of APDs and KVMs from another perspective.

## Acknowledgements

The authors acknowledge financial support from the Australia Research Council under Discovery Project: DP170100363 and Royal Melbourne Hospital Neuroscience Foundation.

